# Methoxsalen inhibited the acquisition of nicotine self-administration: attenuation by cotinine replacement in rats

**DOI:** 10.1101/2023.06.04.543614

**Authors:** Zheng-Ming Ding, Elizabeth M. Neslund, Dongxiao Sun, Xiaoying Tan

## Abstract

Cigarette smoking remains the leading preventable cause of disease and death. Nicotine is the primary reinforcing ingredient in cigarettes sustaining addiction. Cotinine is the major metabolite of nicotine that produces a myriad of neurobehavioral effects. Cotinine supported self-administration and rats with a history of intravenous self-administration of cotinine exhibited relapse-like drug-seeking behavior, suggesting cotinine may also be reinforcing. To date, a potential contribution of cotinine to nicotine reinforcement remains unknown. Nicotine metabolism is mainly catalyzed by hepatic CYP2B1 enzyme in the rat and methoxsalen is a potent CYP2B1 inhibitor. The study tested the hypothesis that methoxsalen inbibits nicotine metabolism and self-administration, and that cotinine replacement attenuates the inhibitory effects of methoxsalen. Acute methoxsalen decreased plasma cotinine levels and increased nicotine levels following subcutaneous nicotine injection. Repeated methoxsalen reduced the acquisition of nicotine self-administration, leading to fewer nicotine infusions, disruption of lever differentiation, smaller total nicotine intake, and lower plasma cotinine levels. On the other hand, methoxsalen did not alter nicotine self-administration during the maintenance phase despite great reduction of plasma cotinine levels. Cotinine replacement by mixing cotinine with nicotine for self-administration dose-dependently increased plasma cotinine levels, counteracted effects of methoxsalen, and enhanced the acquisition of self-administration. Neither basal nor nicotine-induced locomotor activity was altered by methoxsalen. These results indicate that methoxsalen depressed cotinine formation from nicotine and the acquisition of nicotine self-administration, and that replacement of plasma cotinine attenuated the inhibitory effects of methoxsalen, suggesting that cotinine may contribute to the development of nicotine reinforcement.

## Introduction

Tobacco use remains a leading preventable cause of morbidity and mortality. A recent U.S. survey reported that approximately 11.5% of adults were current cigarette smokers and 4.5% adults were current users of e-cigarettes in 2021 ^1^. Nicotine is the main tobacco alkaloid and primary addictive component that reinforces habitual use ^2, 3^. Cotinine is a minor tobacco alkaloid, but the major metabolite of nicotine. It accumulates to much higher plasma levels than nicotine due to its long half-life, making it a widely used biomarker for tobacco use ^4^.

Converging findings from both animal and human research indicate that cotinine is a neuroactive metabolite of nicotine and produces various pharmacological and neurobehavioral effects ^5^. There is evidence suggesting that cotinine may contribute to certain effects of nicotine, e.g., the neuroprotective and cognition-enhancing effects of nicotine ^6–9^. However, potential contribution of cotinine to the addictive effect of nicotine remains unknown.

Several lines of evidence from animal research suggests that cotinine may produce reinforcement-related effects. Cotinine altered food-maintained operant behavior ^10, 11^, and produced nicotine-like discriminative stimulus effects in drug discrimination tests in rodents and non-human primates ^11–13^. Cotinine supported intravenous self-administration in rats and rats with a history of cotinine self-administration exhibited relapse-like drug-seeking behavior ^14, 15^. In addition, cotinine increased brain dopamine transmission in both rodents and non-human primates ^16–20^. Given the importance of dopamine neurotransmission in reinforcment ^21^, these results further suggest cotinine’s role in reinforced behavior.

These findings raise the possibility that cotinine may contribute to the development of nicotine reinforcement. However, testing this hypothesis is not straightforward. One strategy is to block cotinine-specific molecular or cellular mechanisms during nicotine reinforcement. Cotinine is a weak agonist of nicotinic receptors ^22, 23^. Activation of dopamine transmission appeared to be a cellular mechanism underlying cotinine’effects ^19, 20^. However, it is challenging to pharmacologically target these mechanisms without altering nicotine’s effects given both cotinine and niccotine share these mechanisms ^3^. There is evidence indicating a cotinine-specific receptor in the rat brain ^24^. However, lack of characteristics of the receptor and appropriate pharmacological tools makes it premature to target these receptors.

Another strategy is to block cotinine production during nicotine reinforcement. Cotinine formation from nicotine metabolism is mainly catalyzed by cytochrome P450 2A6 enzyme (CYP2A6) in humans, CYP2A5 in mice, and CYP2B1 in rats ^4, 25^. Methoxsalen, a FDA-approved photosensitizer in combination with ultraviolet light for psoriasis treatment, is a potent inhibitor of these enzymes and depresses cotinine formation both *in vivo* and *in vitro* ^26, 27^. If cotinine contributes to the reinforcing effects of nicotine, methoxsalen would be expected to inhibit nicotine reinforcement. Note that methoxsalen also increases plasma nicotine levels ^28^, which may also contribute to the reduced self-administration, confounding the interpretation of the results. To address this challenge, cotinine can be supplemented during methoxsalen treatment to restore plasma cotinine levels. If the effects of methoxsalen is due to reduction of cotinine, cotinine replacement should dampen the inhibitory effect of methoxsalen and increase nicotine self-administration. This provides a feasible approach to investigate the potential role of cotinine in nicotine reinforcement.

The current study examined effects of methoxsalen inhibition of cotinine formation on intravenous nicotine self-adminstration, and then effects of cotinine replacement during methoxsalen treatment to determine its role in nicotine reinforment. The hypothesis is that methoxsalen inhibits cotinine formation and nicotine self-administration, and that cotinine replacement attenuates the inhibitory effects of methoxsalen to increases nicotine self-administration.

## Materials and methods

### Animals

Adult male Wistar rats (starting at ∼ 8 weeks old; Envigo, Indianapolis, IN USA) were housed in a vivarium controlled for temperature and humidity. Our previous study indicates that there was no difference in dose-dependent self-administration of cotinine between male and female rats ^15^. The room was maintained on a reversed 12-h light-dark cycle with light off from 9:30am to 9:30pm. Experimental procedures were performed during the dark phase. Rats were housed in groups (2-4/cage) upon arrival and were acclimated for at least five days. Following surgery, rats were housed individually. Cages were enriched with a polycarbonate play tunnel and nestlets. Food and water were available *ad libitum* unless explicitly stated. Protocols were approved by the Institutional Animal Care and Use Committee at Pennsylvania State University College of Medicine. All experiments were performed in accordance with the principles outlined in the Guide for the Care and Use of Laboratory Animals ^29^.

### Effect of methoxsalen on nicotine metabolism to cotinine

Rats were divided into three groups with each group receiving intraperitoneal injection of one of the solutions: vehicle, 15 mg/kg and 30 mg/kg methoxsalen. The vehicle comprised 95% ethanol, kolliphor EL, and water in a ratio of 1:2:17 in volume. Methoxsalen was dissolved in the vehicle and was injected at a volume of 10 ml/kg. Approximately 30 min later, all rats received a subcutaneous injection of nicotine at 0.5 mg/kg. Nicotine dose was expressed in base throughout the study. Blood samples (∼100 µl) were collected from saphenous vein at 15, 30, 60, 120, 180, 240, 360, and 540 min after the nicotine injection. Blood samples were centrifugated at 14,000 rpm for 20 min at 4°C. Plasma was collected and stored at −80°C.

Plasma levels of nicotine and cotinine were analyzed with liquid chromatography-mass spectrometry as previous described ^30^. Briefly, plasma samples were spiked with deuterated internal standard solution containing nicotine-d3 and cotinine-d3 (Toronto Research Chemicals Inc, Toronto, CA) and acetonitrile/methanol (50/50) solvent. The mixture was centrifugated at 13,000 x *g* for 30 min at 4°C, and the supernatant was loaded onto an Acquity UPLC BEH Hilic column (1.7 µm, 2.1 x 100 mm, Waters, Ireland) in a Sciex EXion UHPLC system. Nicotine and cotinine were separated under a gradient elution condition that started at 15 % solvent A [5 mmol/L NH_4_AC in 50% acetonitrile / 50% water (v/v), pH 6.7] and 85% solvent B [5 mmol/L NH_4_AC in 90% acetonitrile / 10% water (v/v), pH 6.7], and gradually ended at 100% solvent A. Nicotine, cotinine, and internal standards were detected and analyzed with a Sciex QTRAP 6500+ mass spectrometry under the multiple reaction monitoring mode. Signals were quantified by Scie OS 1.5 software.

### Intravenous catheterization and self-administration

Intravenous catheters were implanted following the procedure previously described ^15^. Briefly, rats were anesthetized with 2-3% isoflurane inhalation and the right jugular vein was exposed. Polyurethane tubing [Inner diameter (I.D.) x outer diameter (O.D.) = 0.6 x 1.0 mm; Instech Laboratories, Inc., Plymouth Meeting, PA, USA] of the catheter was inserted into the vein. The remaining portion of the catheter coursed subcutaneously over the shoulder blade to exit the back of the rat via a 22-gauge back-mount cannula (P1 Technologies, Roanoke, VA, USA). Bupivacaine (Hospira, Inc., Lake Forest, IL, USA) at 0.5% and carprofen (Zoetis Inc., Kalamazoo, MI, USA) at 5 mg/kg were applied as analgesia during surgery. Catheters were flushed daily with ∼0.5 ml of 20 IU/ml heparinized saline (Fresenius Kabi, Lake Zurich, IL, USA) containing 0.13 mg/ml gentamicin sulfate (Fresenius Kabi, Lake Zurich, IL, USA). Catheter patency was checked once a week with intravenous administration of ∼0.1 ml of 10 mg/ml methohexital sodium (Par Pharmaceutical, Chestnut Ridge, NY, USA), a fast- and short-acting barbiturate. Rats with failed catheters were excluded from experiments.

Rats recovered for at least 3 days after surgery during which rats were handled on a daily basis. Self-administration was conducted in standard operant chambers (Med Associates Inc., St. Albans, VT, USA) as previously described ^15^. Rats were under light food restriction to maintain at ∼85% body weight. A piece of Froot Loops cereal was placed on the active lever as a bait during the first two sessions to promote exploratory behavior. A fixed-ratio (FR) 1 schedule was employed in which one response on the active lever resulted in an intravenous infusion delivered in 55 µl over a 3-s period via a syringe pump (PHM-100, Med Associates Inc., St. Albans, VT, USA). During infusion, house light was turned off and the cue light above the active lever was turned on. The infusion was followed by a 17-s timeout period during which both cue light and house light were off. Lever presses during the infusion and timeout periods were recorded but produced no further infusions. Responses on the inactive lever were recorded, but no programmed consequence ensued. Sessions were 2 hours in duration and were conducted daily during weekdays.

### Effects of methoxsalen on acquisition and maintenance of nicotine self-administration

Rats were divided into two groups (n = 4/group) trained to self-administer nicotine at 0.03 mg/kg/infusion for 10 sessions. One group received pretreatment of intraperitoneal injection of vehicle and the other received injection of methoxsalen at 15 mg/kg. Pretreatments were given approximately 30 min prior to the beginning of each session. Our previous study indicate that rats typically acquired nicotine self-administration within first 10 sessions ^15^. Immediately following the 10^th^ session, blood samples were collected from saphenous vein and processed for analysis of plasma nicotine and cotinine levels with liquid chromatography-mass spectrometry as described above. Rats were allowed to continue self-administration for additional 6 session without pretreatments to maintain stable self-administration. Then, pretreatments resumed before self-administration sessions. Rats previously receiving vehicle during acquisition sessions were treated with methoxsalen at 15 mg/kg for 8 sessions followed by methoxsalen at 30 mg/kg for 5 sessions. Blood samples were collected at the end of the last session with 15 mg/kg methoxsalen for determination of plasma nicotine and cotinine levels. Rats previously receiving methoxsalen during acquisition sessions were treated with vehicle throughout.

### Effects of cotinine on methoxsalen inhibition of nicotine self-administration

Rats were divided into four groups (n = 4-6/group) and were trained for 10 sessions to acquire self-administration of one of the following nicotine plus cotinine mixture solutions: nicotine remained at 0.03 mg/kg/infusion, and cotinine increased from 0.00375 to 0.0075 to 0.015 and to 0.03 mg/kg/infusion. Prior to each session, rats received intraperitoneal injection of methoxsalen at 15 mg/kg. Then rats continued self-administration with no pretreatment for additional 10 sessions. Blood samples were obtained from saphenous vein at the end of the last treated and non-treated sessions for analysis of plasma nicotine and cotinine levels.

### Effects of methoxsalen on basal and nicotine-stimulated locomotor activity

Locomotor activity was assessed in an open field test following procedures previously described ^31^. Briefly, an open field activity apparatus (Stoelting Co., Wood Dale, IL USA) comprised four opaque walls (L x H: 100 x 35 cm) fit solidly in a nonreflective slotted base. Two opaque quad dividers (L x H: 100 x 35 cm) split the apparatus into four 50 x 50 cm arenas with each arena for one rat. Activities were tracked with the ANY-maze video tracking system (Stoelting Co., Wood Dale, IL USA).

Rats (n=16) were first tested for effects of methoxsalen on basal locomotor activity in 2 test sessions. A cross-over design was employed so that rats received injection of the vehicle in one session and 15 mg/kg methoxsalen in another session. Treatments were randomized and sessions were separated with 3 days. During each session, rats were placed in open field arenas and the injection was conducted at the 30-min mark. Rats were monitored for distance travelled during a 90-min period following the injection. Three rats were excluded from analysis due to loss of tracking during the test.

Rats were then tested for effects of methoxsalen on nicotine-induced locomotor stimulation three days later. Rats were divided into two groups (n = 8 / group) with each group receiving two injections. The first injection was given at the 30-min mark. One group received vehicle injection and the other received methoxsalen at 15 mg/kg. Then at the 60-min mark, all rats received the second injection which was subcutaneous injection of nicotine at 0.5 mg/kg. Rats were monitored for additional 60 min after the nicotine injection for distance travelled. The test procedure was repeated for 9 daily sessions.

### Statistical analysis

Individual data points were expressed as mean ± S.E.M. Plasma nicotine and cotinine levels were analyzed at each time point with one-way ANOVAs or student *t* tests. Number of infusions and lever discrimination during self-administration were analyzed with repeated measures ANOVAs. Significant main effects were followed by post-hoc SNK multiple comparisons. The significant level was set at *p* < 0.05.

## Results

### Methoxsalen depressed nicotine metabolism to cotinine

Plasma nicotine levels decreased whereas cotinine levels increased over time (Fig. 1). Significant differences in blood nicotine levels were observed only at 180, 240, and 360 min after nicotine injection (Fig. 1A; F values > 6.8, *p* values < 0.01). Methoxsalen at both 15 and 30 mg/kg significantly enhanced plasma nicotine levels than the vehicle treatment at these time points. On the other hand, there were significant differences in blood cotinine levels at all time points (Fig. 1B; F values > 18.5, *p* values < 0.001). Methoxsalen at both doses significantly reduced blood cotinine levels compared to the vehicle treatment at all these time points. There was no difference in blood nicotine or cotinine levels between the two doses of methoxsalen.

**Figure 1.**
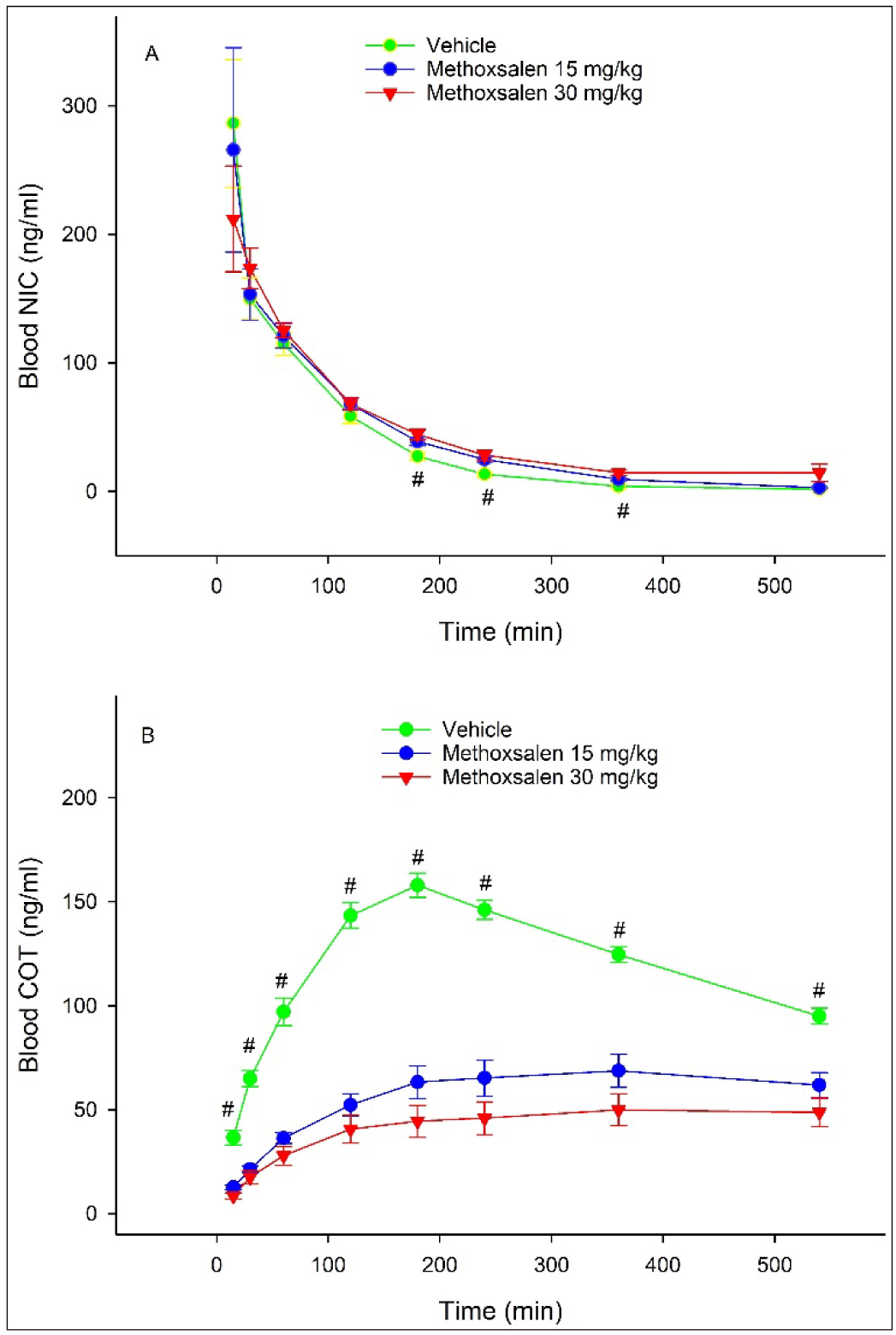
Time-course changes of blood nicotine (A) and cotinine (B) levels following acute injections of methoxsalen (15 & 30 mg/kg, i.p.) and nicotine (0.5 mg/kg, s.c.) in rats. Methoxsalen was injected 30 min prior to the nicotine injection. # *p* < 0.05, significantly different from the methoxsalen treatments.

### Methoxsalen inhibited the acquisition of nicotine self-administration

During acquisition, rats on vehicle, but not on methoxsalen, gradually acquired nicotine infusions (Fig. 2A; session: F_9, 54_ = 3.0, *p* = 0.00; treatment: F_1, 6_ = 9.9, *p* = 0.02; interaction: F_9, 54_ = 8.1, *p* < 0.001). Methoxsalen reduced significantly number of infusions during sessions 3, 4, and 6-9. Methoxsalen also lowered total nicotine intake during acquisition (Fig. 2D; *t*_6_ = 4.7, *p* = 0.003). Although significant in both groups, lever differentiation was more pronounced in vehicle-treated than methoxsalen-treated rats [vehicle (Fig. 2B): session: F_9, 27_ = 2.1, *p* = 0.006; lever: F_1, 3_ = 79.9, *p* = 0.003; interaction: F_9, 27_ = 2.7, *p* = 0.02; Methoxsalen (Fig. 2C): session: F_9, 27_ = 2.8, p = 0.02; lever: F_1, 3_ = 12.0, *p* = 0.04; interaction: F_9, 27_ = 2.5, p = 0.03]. Rats on vehicle elicited more active than inactive responses during sessions 3 throughout 10, whereas rats on methoxsalen showed more on active responses only during sessions 1 and 9. In addition, methoxsalen significantly decreased plasma levels of cotinine (Fig. 2E; *t*_6_ = 5.1, *p* = 0.002), but not nicotine (*t*_6_ = 0.6, *p* = 0.6). These results indicate that methoxsalen reduced cotinine formation and the acquisition of nicotine self-administration.

**Figure 2.**
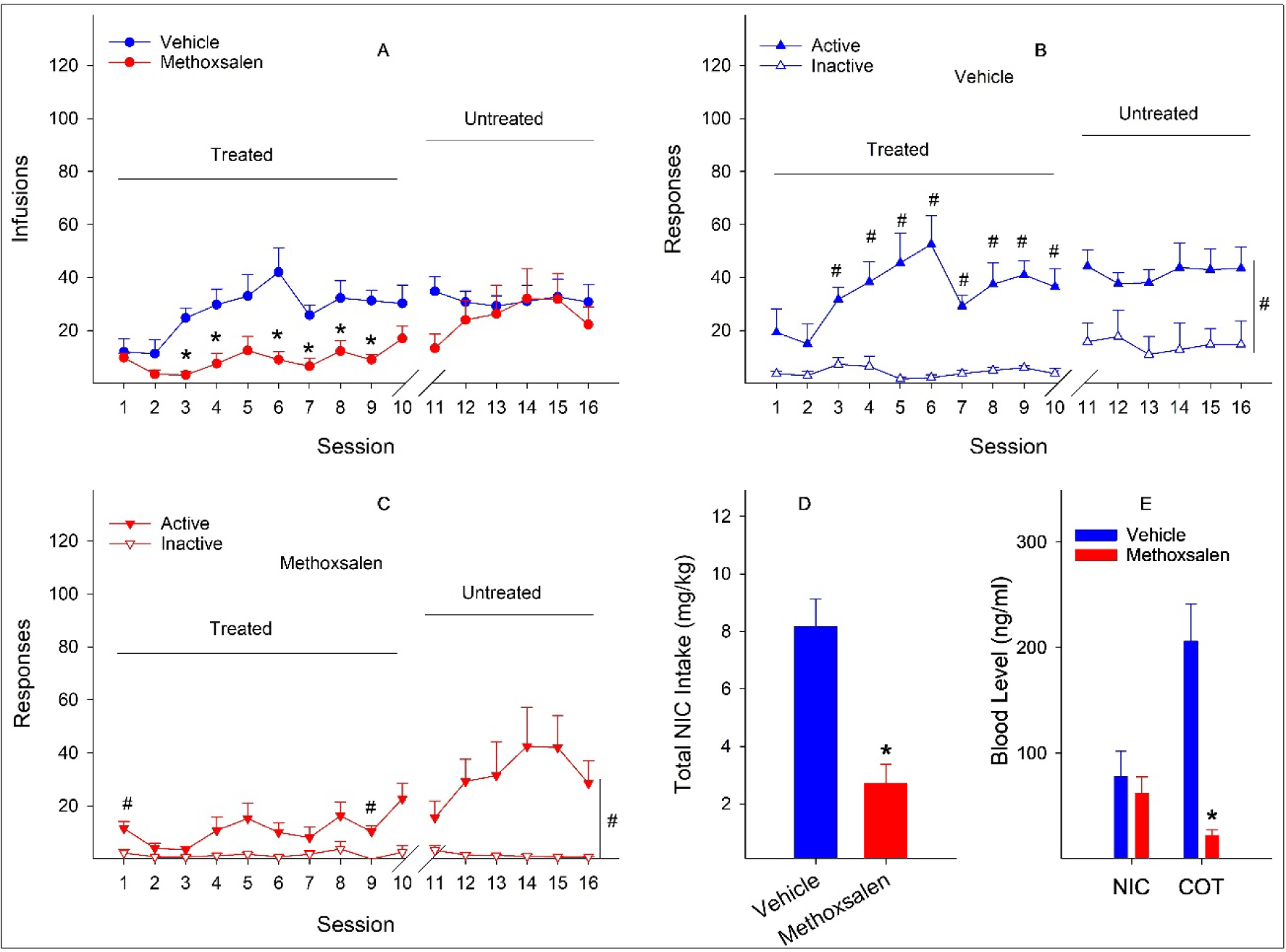
Effects of vehicle and methoxsalen on the acquisition of nicotine self-administration in rats. Methoxsalen (15 mg/kg, i.p.) or vehicle was injected 30 minutes prior to the start of each session. A: Number of infusions per session across treated and untreated sessions. * *p* < 0.05, significantly lower in the methoxsalen-treated group. B & C: Number of responses on active and inactive levers in the vehicle-treated (B) and methoxsalen-treated (C) groups #, *p* < 0.05, significantly greater for active responses. D: Total nicotine intake during the first 10 acquisition sessions. * *p* < 0.05, significantly lower in the methoxsalen-treated group. E: Plasma nicotine and cotinine levels following the 10^th^ acquisition session. * *p* < 0.05, significantly lower in the methoxsalen-treated group.

After the removal of treatments, rats previously treated with methoxsalen gradually increased nicotine infusions to levels similar to those in rats previously treated with vehicle (Fig. 2A; session: F_5, 30_ =1.0, *p* = 0.5; group: F_1, 6_ = 0.6, *p* = 0.5; interaction: F_5, 30_ = 1.6, *p* = 0.2. Rats in both groups maintained lever differentiation (Fig. 2B & 2C).

During maintenance, Methoxsalen did not alter number of nicotine infusions (Fig. 3A; session: F_13, 78_ = 1.9, *p* = 0.04; treatment: F_1, 6_ = 0.04, *p* = 0.95; interaction: F_13, 78_ = 1.4, p = 0.2). The significant effect of session appeared to be due mainly to daily fluctuation of number of infusions. Rats in both groups maintained lever differentiation [vehicle (Fig. 3B): session: F_13, 39_ = 1.3, *p* = 0.3; lever: F_1, 3_ = 42.3, *p* = 0.007; interaction: F_13, 39_ = 1.6, *p* = 0.1; methoxsalen (Fig. 3C): session: F_13, 39_ = 1.9, *p* = 0.6; lever: F_1, 3_ = 231.1, *p* <0.001; interaction: F_13, 39_ = 1.0, *p* = 0.4). Methoxsalen did not alter total nicotine intake (Fig. 3D; *t*_6_ = 0.1, *p* = 0.9). Methoxsalen at 15 mg/kg significantly lowered plasma cotinine (Fig. 3E; *t*_6_ = 4.4, *p* =0.004), but not nicotine levels (Fig. 3E; *t*_6_ = 0.4, *p* = 0.7). These results suggest that methoxsalen did not alter the maintenance of nicotine self-administration despite reduction of cotinine levels.

**Figure 3.**
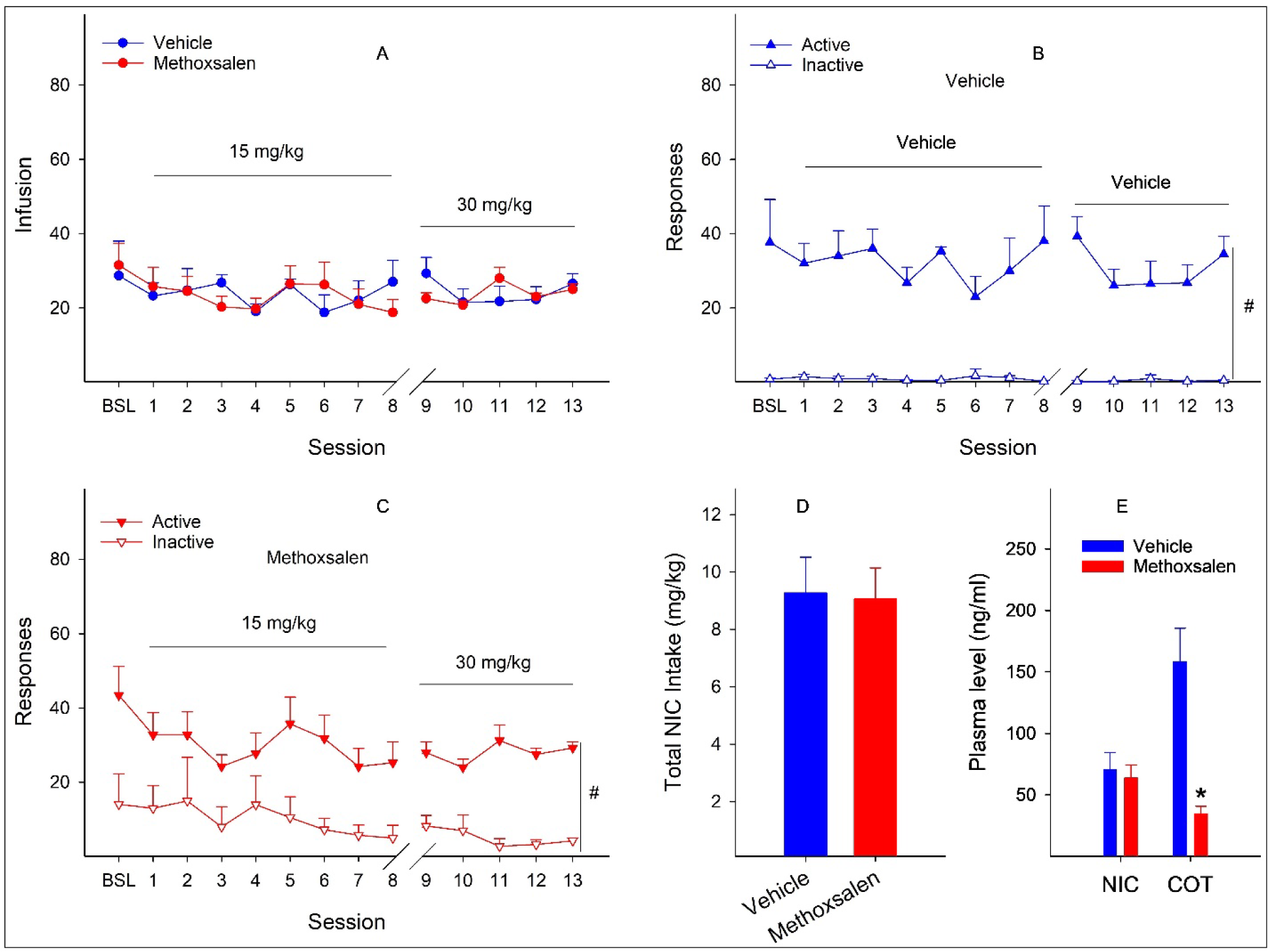
Effects of methoxsalen and vehicle on the maintenance of nicotine self-administration in rats. Methoxsalen (15 & 30 mg/kg, i.p.) or vehicle were injected 30 minutes prior to the start of each session. A: Number of infusions per session. B & C: Number of active and inactive responses in the vehicle-treated (B) and methoxsalen-treated (C) group. #, *p* < 0.05, significantly greater for active responses. D: Total nicotine intake across all sessions. E: Plasma nicotine and cotinine levels following the last session treated with 15 mg/kg methoxsalen. * *p* < 0.05, significantly lower in the methoxsalen-treated group.

### Cotinine replacement dampened methoxsalen inhibition of the acquisition of nicotine self-administration

Mixing cotinine with nicotine for self-administration increased number of infusions in a dose-dependent manner during acquisition (Fig. 4A; session: F_9, 144_ = 16.8, *p* < 0.001; treatment: F_3, 16_ = 5.0, *p* = 0.012; interaction: F_27, 144_ = 1.9, *p* = 0.01). Rats on mixtures with cotinine at 0.015 to 0.03 mg/kg/infusion obtained more infusions than the mixture with cotinine at 0.00375 mg/kg/infusion during sessions 7-10 (Fig. 4A), leading to more nicotine intake in these rats (Fig. 4B; F_3, 16_ = 5.0, *p* = 0.012). Rats on the mixture with cotinine at 0.015 mg/kg/infusion achieved greater cotinine intake than rats on the mixture with cotinine at 0.00375 and 0.0075 mg/kg/infusion, and rats with nicotine plus cotinine at 0.03 mg/kg/infusion obtained more cotinine than all other groups (Fig. 4B; F_3, 16_ = 38.8, *p* < 0.001). Following the 10^th^ acquisition session, plasma samples (Fig. 4C) revealed significant difference in cotinine (F_3, 16_ = 52.4, *p* < 0.001) but not nicotine (F_3, 16_ = 3.3, *p* = 0.05) levels among groups. Rats with nicotine plus cotinine at 0.015 mg/kg/infusion showed greater cotinine levels than rats with nicotine plus cotinine at 0.00375 and 0.0075 mg/kg/infusion. Rats with nicotine plus cotinine at 0.03 mg/kg/infusion obtained greater cotinine levels than all other groups.

**Figure 4.**
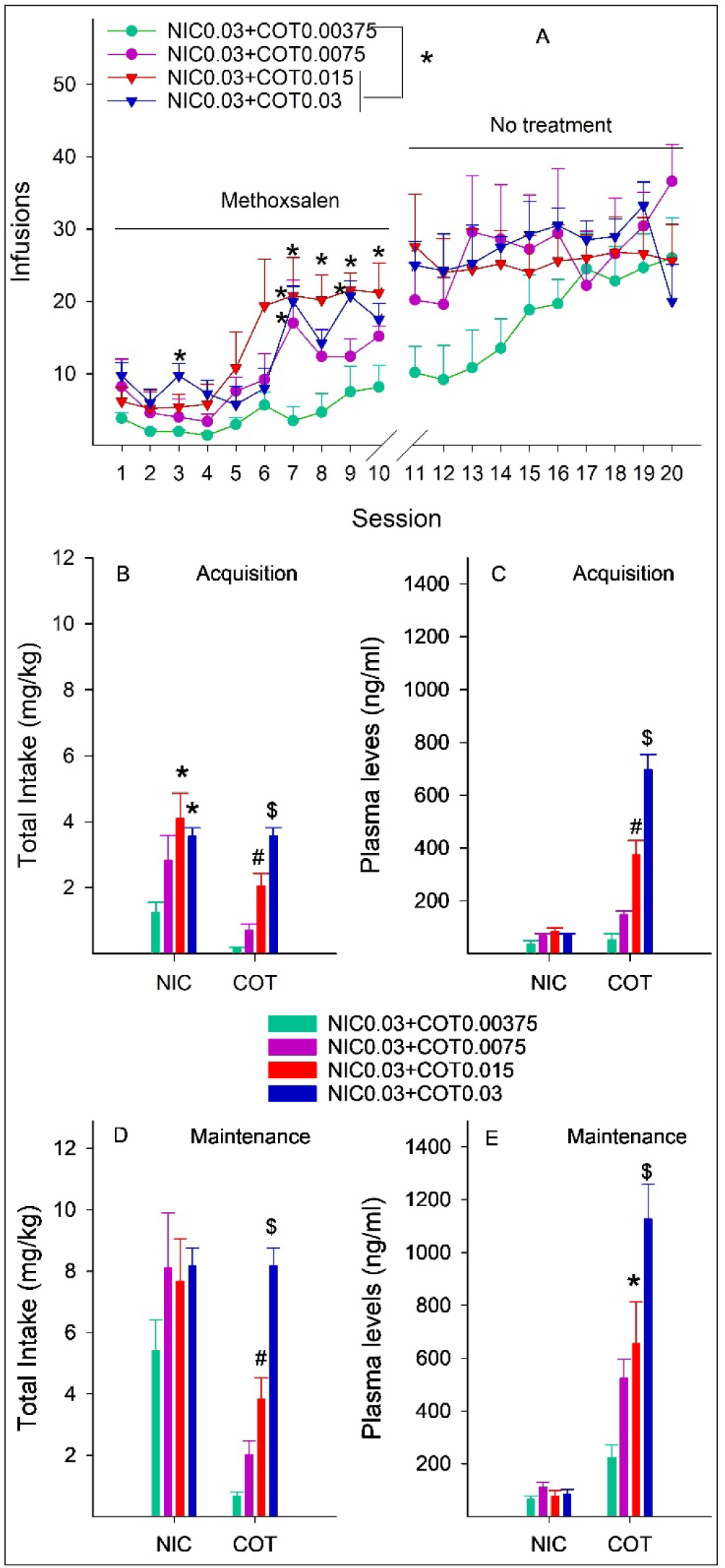
Effects of mixing cotinine (0.00375-0.03 mg/kg/infusion) and nicotine (0.03 mg/kg/infusion) on methoxsalen-inhibited nicotine self-administration in rats. Methoxsalen (15 mg/kg, i.p.) was injected 30 minutes prior to the start of each session. A: Number of infusions per session. * *p* < 0.05, significantly greater than the ‘NIC0.03+COT0.00375’ group. B: Total intake of nicotine and cotinine during the acquisition stage. * *p* < 0.05, significantly greater than the ‘NIC0.03+COT0.00375’ group. # *p* < 0.05, significantly greater than the ‘NIC0.03+COT0.00375’ and ‘NIC0.03+COT0.0075’ groups. $ *p* < 0.05, significantly greater than any other groups. C: Plasma nicotine and cotinine levels after the 10^th^ acquisition session. # *p* < 0.05, significantly greater than ‘NIC0.03+COT0.00375’ and ‘NIC0.03+COT0.0075’ groups. $ *p* < 0.05, significantly greater than any other groups. D: Total intake of nicotine and cotinine during the maintenance stage. # *p* < 0.05, significantly greater than ‘NIC0.03+COT0.00375’ and ‘NIC0.03+COT0.0075’ groups. $ *p* < 0.05, significantly greater than any other groups. E: Plasma nicotine and cotinine levels after the 10^th^ acquisition session. * *p* < 0.05, significantly greater than the ‘NIC0.03+COT0.00375’ group. $ *p* < 0.05, significantly greater than any other groups.

After discontinuation of methoxsalen, rats continued to increase infusions with no significant difference among groups (Fig. 4A; session: F_9, 144_ = 3.3, *p* = 0.001; group: F_3, 16_ = 1.1, *p* = 0.4; interaction: F_27, 144_ = 1.7, *p* = 0.032). No significant difference during any individual session was revealed in post-hoc analyses, suggesting that the significant effect of interaction was mainly driven by the significant effect of session. Rats with cotinine at 0.015 mg/kg/infusion showed greater cotinine intake than rats with cotinine at 0.00375 and 0.0075 mg/kg/infusion, and rats with cotinine at 0.03 mg/kg/infusion obtained more cotinine intake than all other groups (Fig. 4D; F_3, 16_ = 7.8, *p* = 0.002). There was no significant difference in nicotine intake (Fig. 4D; F_3, 16_ = 1.2, *p* = 0.4). Following the last session, there was significant difference in plasma cotinine (F_3, 16_ = 43.6, *p* < 0.001), but not nicotine (F_3, 16_ = 1.4, *p* = 0.05) levels (Fig. 4E. Rats with cotinine at 0.015 mg/kg/infusion showed greater cotinine levels than rats with cotinine at 0.00375 mg/kg/infusion. Rats with cotinine at 0.03 mg/kg/infusion achieved higher cotinine levels than all other groups.

### Methoxsalen had no effect on basal and nicotine-stimulated locomotor activity

At basal state (Fig. 5A), there was no significant difference in distance travelled between groups (F_1, 12_ = 1.9, *p* = 0.2), suggesting that methoxsalen did not alter basal locomotor activity.

**Figure 5.**
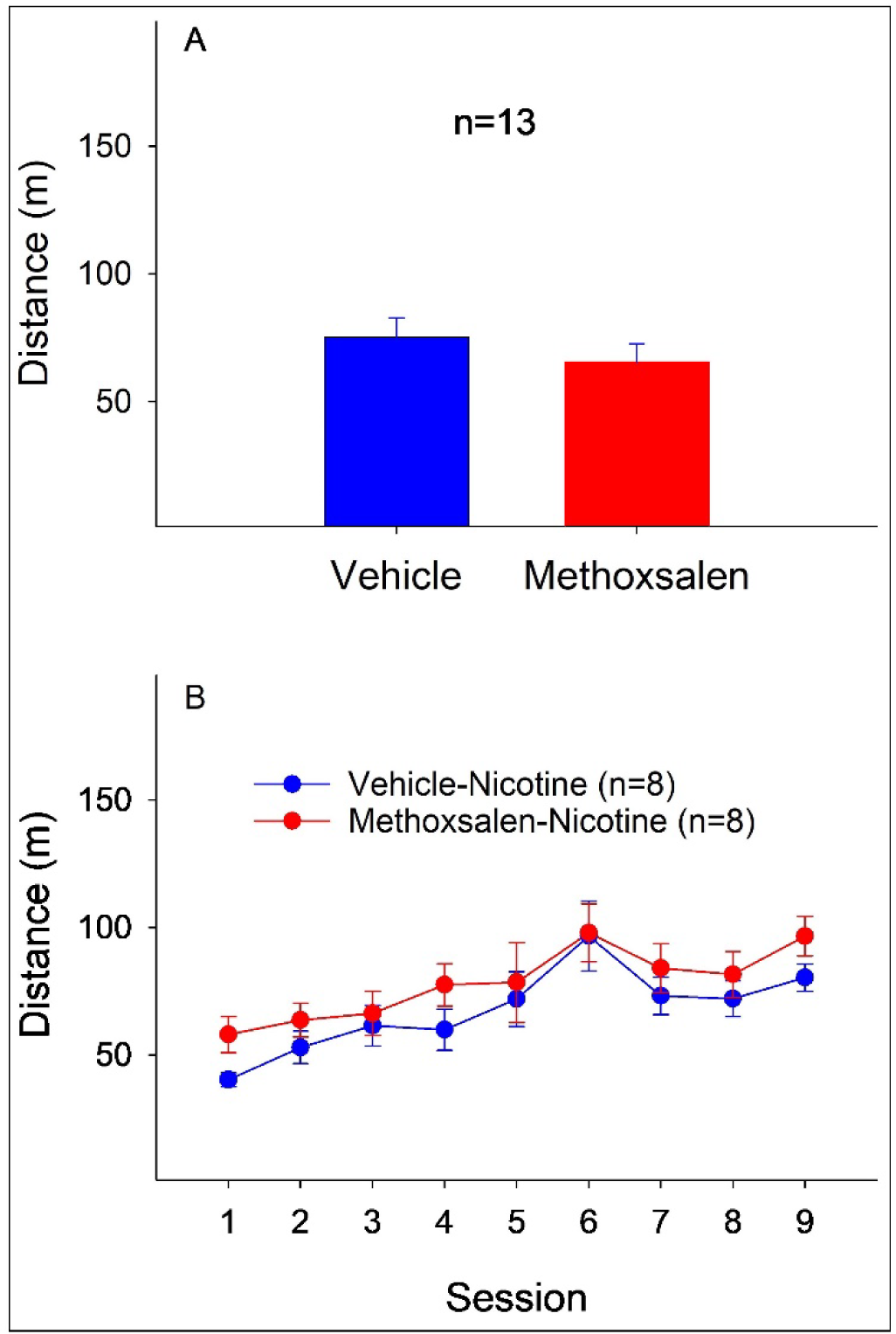
Effects of vehicle and methoxsalen on basal (A) and nicotine-induced (B) locomotor activity in the open field test.

For nicotine-induced locomotor stimulation, there was no significant effect of treatment (F_1, 14_ = 1.2, *p* = 0.3) or the session x treatment interaction (F_8, 112_ = 0.4, *p* = 0.9). However, a significant effect of session (F_8, 112_ = 11.4, *p* < 0.001) was observed due to gradual increase of distance travelled during repeated nicotine injection, suggesting the development of locomotor sensitization. These results suggest that methoxsalen did not change nicotine-induced locomotor stimulation or sensitization.

## Discussion

The current study found that methoxsalen inhibited nicotine metabolism following acute subcutaneous nicotine injection, resulting in decreased plasma cotinine levels and increased nicotine levels. Repeated methoxsalen reduced, in a reversible manner, the acquisition of nicotine self-administration, leading to fewer infusions and lack of lever differentiation during most acquisition sessions, as well as lower total nicotine intake and plasma cotinine levels compared to repeated vehicle treatments. Methoxsalen did not alter the maintenance of nicotine self-administration despite significant reduction of plasma cotinine levels. In the presence of methoxsalen, rats on mixtures of cotinine and nicotine exhibited dose-dependent increase of acquisition of self-administration, number of infusions, total intake of nicotine and cotinine, and plasma levels of cotinine. Of further note, methoxsalen did not alter basal locomotor activity or nicotine-induced locomotor stimulation and sensitization. These results indicate that inhibition of nicotine metabolism decreased the development of nicotine self-administration, and concurrent administration of cotinine and nicotine enhanced the development of self-administration despite the inhibition of nicotine metabolism. These results suggest that cotinine may contribute to the development of nicotine reinforcement.

Nicotine conversion to cotinine in the rat is preferentially catalyzed by CYP2B1, which accounts for ∼40% of nicotine metabolism *in vitro* ^25, 32^. Methoxsalen is a potent CYP2B1 inhibitor with Ki value at 2.9 µM ^26^, and has been shown to inhibit 58% of nicotine metabolism in rat liver microsomes ^33^. As a result, methoxsalen in the current study reduced plasma cotinine levels and increased plasma nicotine levels *in vivo* after subcutaneous nicotine injection. These results are consistent with previous findings in humans and mice. Nicotine is mainly metabolized by CYP2A6 in humans and by CYP2A5 in mice, and these two enzymes are orthologous and share 84% amino acid homology ^4, 34, 35^. Methoxsalen inhibits these enzymes with its potency at 0.1-5.4 µM for CYP2A6 ^36–38^ and 0.32-1 µM for CYP2A5 ^28, 36^. In acutely abstinent cigarette smokers, methoxsalen increased plasma nicotine levels and decreased nicotine clearance following nicotine administration ^38, 39^. In mice, methoxsalen enhanced plasma levels and half-life of nicotine, and decreased plasma cotinine following nicotine administration ^28, 40^.

Effects of methoxsalen on nicotine metabolism seem to be species-dependent. Methoxsalen-induced increase of nicotine levels occurred as late as 180 min in rats with ∼60-100% increase (Fig. 1), in contrast to 5-15 min in mice with 150-200% increase ^28, 40^ and 30-60 min in humans with 40-100% increase ^38, 39^. In addition, maximal nicotine levels were not altered by methoxsalen in rats (Fig. 1), but increased in mice ^40, 41^ and humans ^38, 39^. The mechanism responsible for these species-dependent differences remains unknown. It is noted that these species show differences in effectiveness of nicotine metabolism. CYP2A6 is an effective enzyme and metabolizes approximately 80% of nicotine in humans ^42, 43^. CYP2A5 in mice is even more efficient than CYP2A6 in humans ^28, 34, 35^. In rats, the CYP2B1 only accounts for an average of ∼50% of nicotine metabolism ^25, 33^. Whether these differences contribute to the observed difference in methoxsalen on nicotine remains to be determined.

Repeated methoxsalen reduced nicotine self-administration during the acquisition, but not maintenance, phase (Fig. 2 & 3). In addition, methoxsalen did not alter locomotor activity at basal state and during nicotine treatment (Fig. 5), suggesting that effects of methoxsalen on self-administration were unlikely due to nonspecific locomotor inhibition. These results further suggest that the inhibition of nicotine metabolism may regulate the development of nicotine reinforcement in rats. A previous study reported that a novel CYP2A6 inhibitor, DLCI-1, reduced nicotine self-administration during the maintenance phase in mice ^44^. On the other hand, methoxsalen was found to have minimum effect on human smoking ^38, 39^. It remains to be determined whether methoxsalen may alter acquisition of nicotine use in mice and humans. However, the difference in methoxsalen’s effects on maintenance of nicotine use may reflect a species-specific effect. Interestingly, concomitant administration of methoxsalen with nicotine decreased the desire to smoke, increased the interval between cigarettes, and reduced total number of cigarette smoked in smokers ^38^, suggesting that methoxsalen may be beneficial as a combination therapy with nicotine.

It is noted that methoxsalen did not alter plasma nicotine levels during active self-administration (Fig. 2 & 3). Two factors may have potentially contributed to the lack of effect. First, our study indicates that methoxsalen-induced nicotine increase occurred as late as 180 min (Fig. 1), which is significantly greater than the duration of the session at 120 min. A second potential factor is the dynamic process of nicotine intake, metabolism and clearance during the session, making it difficult, if not impossible, to detect difference in plasma nicotine levels. Similar results were reported in human studies in which methoxsalen did not alter plasma nicotine or cotinine levels over a 3-day regimen with blood samples compared before and after the treatment ^39^. The Chen et al study (2020) did not report plasma nicotine levels, therefore it remains unknown how DLCI-1 may change nicotine levels during active self-administration in mice.

To test a potential role of cotinine in nicotine reinforcement, the methoxsalen-reduced plasma cotinine levels were restored by mixing cotinine with nicotine solution for self-administration. This resulted in dose-dependent increase of plasma cotinine levels, which led to dose-dependent enhancement of acquisition of self-administration despite the methoxsalen treatment (Fig. 4). These results indicate that restoration of plasma cotinine levels dampened the inhibitory effects of methoxsalen, suggesting that the inhibitory effect of methoxsalen on the acquisition of nicotine self-administration was, at least in part, due to reduction of plasma cotinine levels.

These results suggest that cotinine may facilitate the acquisition of nicotine self-administration, thus contributing to the development of nicotine reinforcement. It should be noted that cotinine and nicotine co-self-administration results in rapid delivery of high levels of cotinine to the circulating system, which is conceivably different from the slow dynamic process of cotinine formation and distribution through nicotine metabolism. Therefore, the definitive role of cotinine on nicotine reinforcement in natural state should be further tested in the future when new technology emerges that allows sequestering cotinine without altering the dynamic of nicotine metabolism and cotinine formation, e.g., agents that may facilitate cotinine renal clearance or block cotinine entry into the brain to prevent its neurobehavioral effects.

Note that studies from mice indicate that reduced nicotine metabolism through either methoxsalen or CYP2A(4/5) null mice resulted in increased nicotine-induced conditioned place preference (CPP), suggesting that reduction of cotinine levels may be associated with an increase in the rewarding effects of nicotine ^41, 45^. This difference may reflect a species-specific effect. This may also be paradigm-specific, i. e., experimenter-administered nicotine in CPP vs animal-controlled active intake of nicotine during self-administration. In addition, it is possible that the enhancing effect from increased nicotine as a result of decreased metabolism may have outweighed the attenuating effects from reduced cotinine, leading to a net increase of CPP. This possibility may be tested in the future by examining effect of cotinine restoration after methoxsalen on CPP. If it further increases CPP, that will suggest that cotinine may contribute to the development of CPP.

Our findings suggest that nicotine metabolism plays an important role in the development of nicotine reinforcement in rats. Human studies indicate that individuals with genetic deficiencies, e.g., CYP2A6 deletion allele (CYP2A6*4), are associated with lower risk of becoming habitual smokers ^46, 47^. CYP2A6*4 leads to substantial reduction in nicotine metabolism and lower levels of plasma cotinine levels after smoking or nicotine administration ^4, 48^. The protective effects of these genetic deficiencies have been mainly attributed to increased nicotine bioavailability, prolonged nicotine actions, and reduced nicotine clearance. Our findings suggest that reduced cotinine formation may impair development of nicotine reinforcement, which may contribute to the protective effects of CYP2A6 genetic deficiencies. Given this, it will be interesting to examine whether cotinine may contribute to effects of methoxsalen or DLCI in smokers and mice, respectively ^38, 39, 44^. In addition, methoxsalen also altered other nicotine-induced pharmacological effects, e.g., antinociception, hypothermia ^28, 40^. It would be interesting to determine in future studies whether cotinine may contribute to these effects of nicotine.

In summary, our study reveals that in rats methoxsalen reduced the acquisition of nicotine self-administration, which was associated with reduced plasma cotinine levels, and that restoration of plasma cotinine during nicotine self-administration attenuated the inhibitory effects of methoxsalen to enhance the acquisition of nicotine self-administration. These results suggest that cotinine may facilitate the development of nicotine reinforcement. In addition, the current approach may be adapted to examine the role of other nicotine metabolites in the development of nicotine reinforcement, e.g., nornicotine which derives from nicotine metabolism by CYP2A6 and produces its own reinforcing effects in rats ^34, 49^.

## Acknowledgements

This study was supported in part by the US National Institutes of Health grant DA044242. The authors declare no conflict of interest. The content of this manuscript is solely the responsibility of the authors and does not necessarily represent the official views of National Institutes of Health.

## Authorship Contributions

Research conceptualization and design: Ding

Data collection: Neslund, Tan, Sun

Data analysis: Ding, Neslund, Tan, Sun

Manuscript writing/review/editing: Ding, Neslund, Tan, Sun

